# Transcriptome signatures of the medial prefrontal cortex underlying GABAergic control of resilience to chronic stress exposure

**DOI:** 10.1101/2024.07.10.602959

**Authors:** Meiyu Shao, Julia Botvinov, Deepro Banerjee, Santhosh Girirajan, Bernhard Lüscher

## Abstract

Analyses of postmortem human brains and preclinical studies of rodents have identified somatostatin (SST)-positive interneurons as key elements that regulate the vulnerability to stress-related psychiatric disorders. Conversely, genetically induced disinhibition of SST neurons or brain region-specific chemogenetic activation of SST neurons in mice results in stress resilience. Here, we used RNA sequencing of mice with disinhibited SST neurons to characterize the transcriptome changes underlying GABAergic control of stress resilience. We found that stress resilience of male but not female mice with disinhibited SST neurons is characterized by resilience to chronic stress-induced transcriptome changes in the medial prefrontal cortex. Interestingly, the transcriptome of non-stressed stress-resilient male mice resembled the transcriptome of chronic stress-exposed stress-vulnerable mice. However, the behavior and the serum corticosterone levels of non-stressed stress-resilient mice showed no signs of physiological stress. Most strikingly, chronic stress exposure of stress-resilient mice was associated with an almost complete reversal of their chronic stress-like transcriptome signature, along with pathway changes indicating stress-induced enhancement of mRNA translation. Behaviorally, the mice with disinhibited SST neurons were not only resilient to chronic stress-induced anhedonia — they also showed an inversed anxiolytic-like response to chronic stress exposure that mirrored the chronic stress-induced reversal of the chronic stress-like transcriptome signature. We conclude that GABAergic dendritic inhibition by SST neurons exerts bidirectional control over behavioral vulnerability and resilience to chronic stress exposure that is mirrored in bidirectional changes in expression of putative stress resilience genes, through a sex-specific brain substrate.

## INTRODUCTION

Chronic and excessive amounts of stress are vulnerability and symptoms-precipitating factors for virtually all psychiatric disorders, especially depressive disorders, posttraumatic stress disorder (PTSD), and schizophrenia (SCZ). However, individuals differ significantly in their susceptibility to stress, pointing to differences in stress resilience, a feature that has been described by the American Psychological Association as “the process and outcome of successfully adapting to difficult or challenging life experiences” (https://dictionary.apa.org/resilience, last checked June 6, 2024). Clinical and preclinical studies have identified somatostatin (SST)-positive GABAergic interneurons in the frontal cortex as key elements regulating the vulnerability to stress. Specifically, SST protein and mRNA and other transcripts that map to these neurons are downregulated in postmortem brain of subjects who died with major depressive disorder (MDD), bipolar disorder (BP), or SCZ, as well as in association with aging and Alzheimer’s disease ^1–7^. Preclinical studies suggest that reduced SST neuron function may causally contribute to these conditions, as SST is downregulated following chronic stress exposure ^8^ and deliberately inhibiting SST neuron activity leads to heightened emotional behavior and cognitive deficits associated with mental disorders and aging ^9–11^.

SST interneurons preferentially innervate the distal dendrites of pyramidal cells, modulating the strength of excitatory inputs ^12, 13^. Feedforward inhibition mediated by SST neurons scales with the strength of their excitatory input from the basolateral amygdala ^14^. In addition, inhibitory synaptic inputs from SST neurons onto pyramidal cell dendrites are strengthened post-synaptically by hetero-synaptic NMDA-receptor-mediated plasticity ^15^. Together, these features predict that GABAergic inhibition of pyramidal cell dendrites by SST neurons exerts a naturally neuroprotective role that is amplified during increased network load but may get compromised under psychopathological and chronic stress conditions, as well as during aging ^7, 11^. Indeed, a single 1-h immobility stressor leads to lasting activation of SST neurons ^14^. Consistent with the neuroprotective role of SST neurons, we showed that mice with globally disinhibited SST neurons (through cell type-specific inactivation of the *Gabrg2* gene in SSTCre:γ2^f/f^ mice) exhibit biochemical and behavioral alterations that mimic the effects of antidepressant drug treatment, including resilience to the anxiogenic effects of uncontrolled chronic mild stress exposure ^8, 16^. Therefore, increased excitability of SST neurons in SSTCre:γ2^f/f^ mice strengthens naturally stress-induced neuroprotective circuits that lead to stress resilience. Here, we adopted the Chronic Variable Stress (CVS) paradigm ^17, 18^ to further characterize the resiliency phenotype of SSTCre:γ2^f/f^ mice under more severe chronic stress conditions and to elucidate transcriptome signatures and mechanisms that underlie GABAergic induction of stress resilience.

In a landmark study elucidating gene expression changes associated with stress resilience in the chronic social defeat stress (CSDS) model, Krishnan et al. showed that resilience is an active process involving significantly more stress-induced gene expression changes than observed in stress-vulnerable mice ^19, 20^. A strength of this model is that it makes no prior assumptions regarding circuits and gene expression changes that mediate stress resilience. The genetically induced SSTCre:γ2^f/f^ model used here differs in that it focuses a priori on GABAergic microcircuits that promote stress resilience via cortical brain regions ^8, 16, 21^. This model comes with the key advantage that it allows for the molecular comparison of non-stressed (NS) stress-resilient mice with stressed or NS stress-susceptible mice as well as stressed stress-resilient mice — a feature that is not possible with the CSDS model. Therefore, the SSTCre:γ2^f/f^ model allows testing whether stress-resilient mice differ from stress-vulnerable mice with respect to physiological parameters that do not involve chronic stress exposure. Lastly, unlike the standard CSDS model, the SSTCre:γ2^f/f^ model is readily amenable to both sexes.

CVS exposure of SSTCre:γ2^f/f^ mice revealed resilience to CVS with respect to both anxiety- and anhedonia-related changes in motivated behavior in both sexes. However, RNA sequencing (RNA-Seq) of the mPFC revealed that SST neuron-mediated resilience in this brain region is male-specific. Focusing on male mice, we then compared the CVS-induced transcriptome changes of stress-vulnerable mice with those of stress-resilient mice, as well as with transcriptome changes between NS stress-vulnerable and stress-resilient mice. We found that stress resilience is associated with stress-induced upregulation of mRNA translation while stress vulnerability is associated with downregulation of mRNA translation, cell adhesion and diverse inter- and intracellular signaling pathways. Remarkably, the transcriptome changes of NS stress-resilient mice partly mimicked the transcriptome signature of CVS-exposed stress-vulnerable mice. Most strikingly, CVS exposure of stress-resilient mice resulted in the reversal of the transcriptome changes of NS stress-resilient vs NS stress-vulnerable mice.

## MATERIALS AND METHODS

### Animals

All animal experiments were approved by the Institutional Animal Care and Use Committees (IACUC) of The Pennsylvania State University and performed in accordance with guidelines of the National Institutes of Health (NIH). SSTCre mice (also known as Sst ^tm2.1(cre)Zjh/J^, Stock No. 013044) and C57BL/6J mice (BL6, Stock No. 000664) were obtained from Jackson Laboratory (Bar Harbor, ME, USA). The γ2^f/f^ mouse line (*Gabrg2^tm2Lusc^*/J, Stock No: 016830, Jackson Laboratory) containing a *Gabrg2* allele flanked by loxP sites was generated in house^22^. All mice were backcrossed to the BL6 strain for at least five generations and maintained on a 12:12 h normal light-dark cycle with food and water available *ad libitum* on corn cob bedding. The mice were genotyped by PCR of tail DNA at the time of weaning as described on the JAX website, separated by genotype and sex into experimental groups, and then moved to a 12:12h reversed light-dark cycle. Male and female experimental mice were maintained on separate cage racks and, whenever possible, in separate rooms to inhibit the estrus cycling of females ^23^. All experiments were done with the experimenter blinded to genotype and treatment.

### Chronic variable stress treatment

Experimental groups of mice that differed in sex and genotype were further divided into NS and CVS groups with balancing for sucrose preference and body weight and housed 2-3 per cage. CVS exposure was initiated at the age of 8–10 weeks and included three different stressors repeated for a total of 21 days ^17, 18^, starting on day one with a 1-h tail suspension stressor, followed on day two with a 1-h restraint stressor during which the mice were placed into perforated 50 ml falcon tube, and followed on day three with exposure to 100 randomly distributed foot shocks (0.45 mA x 3 s) within 1 h, using max 10 mice per chamber of a two-compartment shuttle box with the connecting gate closed (SanDiego Instruments, San Diego, CA). After each stressor, the mice were returned to their home cage. At the end of CVS treatment, all mice were singly housed in preparation for further analyses.

### Behavioral analyses

Behavioral tests were initiated 18–24 h after the last stressor by an experimenter blinded to genotype and CVS treatment, starting 1 h after the lights went off. All behavioral experiments (except for the open field test (OFT)) were conducted under red light, starting with the novelty suppressed feeding test (NSFT), followed by the sucrose splash test (SSPT), sucrose preference test (SPT), female urine sniffing test (FUST) for males only, and OFT. For the NSFT ^16, 24^, the mice were food-deprived for 18 h, transferred to the corner of a novel Plexiglass arena (50 x 50 x 20 cm) containing three cm of saw dust bedding and a pellet of rodent chow placed on a white cotton nesting square (6 x 6 x .5 cm) in the center of the arena. The latency to feed was hand-scored with feeding defined as the mouse biting into the chow while resting on its hind paws. Trials were stopped after 5 min, even if no feeding occurred. For the OFT, the mice were allowed to explore an odor-saturated arena of 50 x 50 x 20 cm with opaque Plexiglass walls and a transparent floor underlaid with white reflective paper, exposed to white light (75 lux). The distance traveled within the first 5-min was recorded using Ethovision XT (Noldus Information Technologies, Leesburg, VA). For the SSPT ^25^, the mice were transferred to an empty cage and sprayed on their backs with 1 mL of 10% sucrose solution to stimulate grooming behavior. The mice were immediately returned to their home cage and the grooming duration was scored manually for 5 min. For the SPT ^17^ the mice were trained to drink water from two 25 mL sealed plastic pipettes for 24 h. The following day, one of the pipettes was replaced with 1% sucrose for 24 h. The pipettes were weighed at the start and end of the trial. Sucrose preference was defined as the ratio of sucrose consumption to the total liquid consumption. For the FUST ^26^, male mice were accustomed to a sterile cotton-tip applicator (Patterson Veterinary Supply, Saint Paul, MN) in their home cage for 30 min. For the test, the mice were first exposed to a new water-soaked cotton-tip for 3 min. After 45 min they were exposed to a fresh female-urine-soaked cotton-tip for another 3 min. The behavior was video-recorded, and the duration spent sniffing the female urine was scored offline. Notably, the cages did not have cage lids or wire tops during the entire procedure, and the test is applicable to male mice only.

### Analyses of gene expression by RNA-Seq

#### Library preparation and RNA-Seq

Mice were sacrificed by cervical dislocation 24 hours after the last stressor. The mPFC was dissected from 1 mm coronal sections prepared using a mouse brain matrix (Stoelting Co., Wood Dale, IL), followed by extraction of total RNA using a GenElute™ Mammalian Total RNA Miniprep Kit and on-column DNase I treatment (Qiagen). The RNA integrity number and concentration were assessed using a Fragment Analyzer (Agilent 5400). Messenger RNA was purified from 400 ng of total RNA, and libraries were constructed using the NEBNext Ultra™ II Directional RNA Library Prep Kit for Illumina (New England Biolabs). The libraries were sequenced on an Illumina NovaSeq 6000 system at a depth of 20 million 150-bp paired-end reads per sample. RNA reads were filtered using fastp v.0.23.2 ^27^ to remove sequencing adapters and reads shorter than 50-bp and aligned to the mouse genome GRCm39 using STAR aligner v.2.7.10b ^28^. The STAR genome index was constructed using the annotation v.M32 downloaded from GENCODE. FeatureCounts ^29^ was utilized to quantify read pairs aligned to exons at the gene level in a strand-specific mode. Read pairs aligned to multiple genes were excluded from the analysis. The sequence data can be accessed from NCBI (Accession Number)

#### Differential Expression Analysis

mRNA read pair counts were extracted from RNA-Seq data and differential expression analysis was performed using DESeq2 v.1.40.2 ^30^. Genes were selected for further analysis if they had more than 5 reads in a number of samples equal to or exceeding the size of the smallest experimental group. A cutoff of p < 0.01 was used to identify differentially expressed genes (DEGs). The correlation coefficients of Log2 fold changes (FC) between different contrasts were computed using the Spearman method. The lists of DEGs are available in **Tables S1–S11**.

#### Subsampling and Subsequential Differential Expression Analysis

Three samples were randomly selected from each condition, considering all possible combinations. Differential expression analyses were then conducted for all possible subsampled sets of samples to generate multiple DEG counts for each of the contrasts compared. If the number of sample combinations varied between contrasts, a random selection of samples was performed to equalize the sample numbers across contrasts.

#### Principal Component Analysis (PCA)

PCA was conducted using the DESeq2 R package ^30^. Batch differences between male and female samples were removed using ComBat-seq ^31^ followed by a variance stabilizing transformation. The top 1,000 genes based on variance were used for PCA using the plotPCA function.

#### Pathway enrichment analysis

Pathway enrichment analyses were conducted using Ingenuity Pathway Analysis (IPA) (QIAGEN, Inc.). A cutoff of p < 0.05 was applied to select altered pathways. The pathway activation and inhibition states were assessed using IPA’s activation Z-score tool. Pathway comparisons were done using IPA’s integrated comparison analysis tool and ranked by Z-score. For clarity, the pathways related to coronavirus pathogenesis, cancer, autism and pancreatic secretion were excluded from the rankings. Pathway lists are available in **Tables S12–S17**.

#### Disease enrichment analysis

Disease enrichment analysis was performed on Enrichr (https://maayanlab.cloud/Enrichr/) ^32–34^ utilizing the DisGeNET library ^35–38^ after the mouse gene identifiers (IDs) were transferred to human gene IDs using the SynGO ID convert tool ^39^. A cutoff of p < 0.05 was applied to identify enriched diseases. Genes associated with the enrichment terms MDD (sum of MDD, unipolar depression and depressive disorders), BP, PTSD, and SCZ were extracted for further analysis.

### Measurement of serum corticosterone

Serum corticosterone (CORT) was measured nine days after the end of CVS, five to seven hours after the start of the dark phase, using an ELISA kit (Enzo Life Sciences, Farmingdale, NY) and a SepctraMax® i3x microplate reader (Molecular Devices LLC, San Jose, CA) and SoftMax® Pro 7 Software (Molecular Devices LLC).

### Statistical analyses

Statistical analyses were performed using Prism 10 software (GraphPad, La Jolla, CA). Outliers identified with the ROUT method were omitted from analyses. Pairwise comparisons of data that satisfied the normality assumption were compared using a two-tailed Student’s t-test. Data that failed the equal variance assumption were compared using Welch’s two-sided t-test. Two- and three-way ANOVAs were employed to analyze multi-group means with Tukey’s post hoc testing. Data that did not meet the normality assumption were log transformed before further analyses or analyzed using Mann Whitney tests.

## RESULTS

### SSTCre:γ2^f/f^ mice are resilient to CVS-induced changes in motivated behavior independent of sex

We previously reported that SSTCre:γ2^f/f^ mice are resilient to the anxiogenic effects of uncontrolled chronic mild stress exposure ^8^. However, this protocol had failed to reliably induce anhedonia-like changes in rewarding behavior and thereby prevented us from testing the mice for resilience in this behavioral domain. Here we adopted a CVS protocol (Figure 1A), which in SSTCre control mice results in both anxiety-like and anhedonia-like changes in motivated behavior. Weekly measurements of body weight during CVS exposure revealed similar stress-induced attenuation of body weight gain in SSTCre and SSTCre:γ2^f/f^ male and female mice (**Figure 1B, F**). Thus, with respect to whole body physiology all mice seemed to experience stress similarly, independent of genotype and sex. Separate cohorts of SSTCre, SSTCre:γ2^f/+^, SSTCre:γ2^f/f^ mice were then subjected to CVS or NS control conditions to test for stress-induced changes in negatively (NSFT) and positively regulated motivated behavior (FUST, SSPT, SPT).

**Figure 1.**
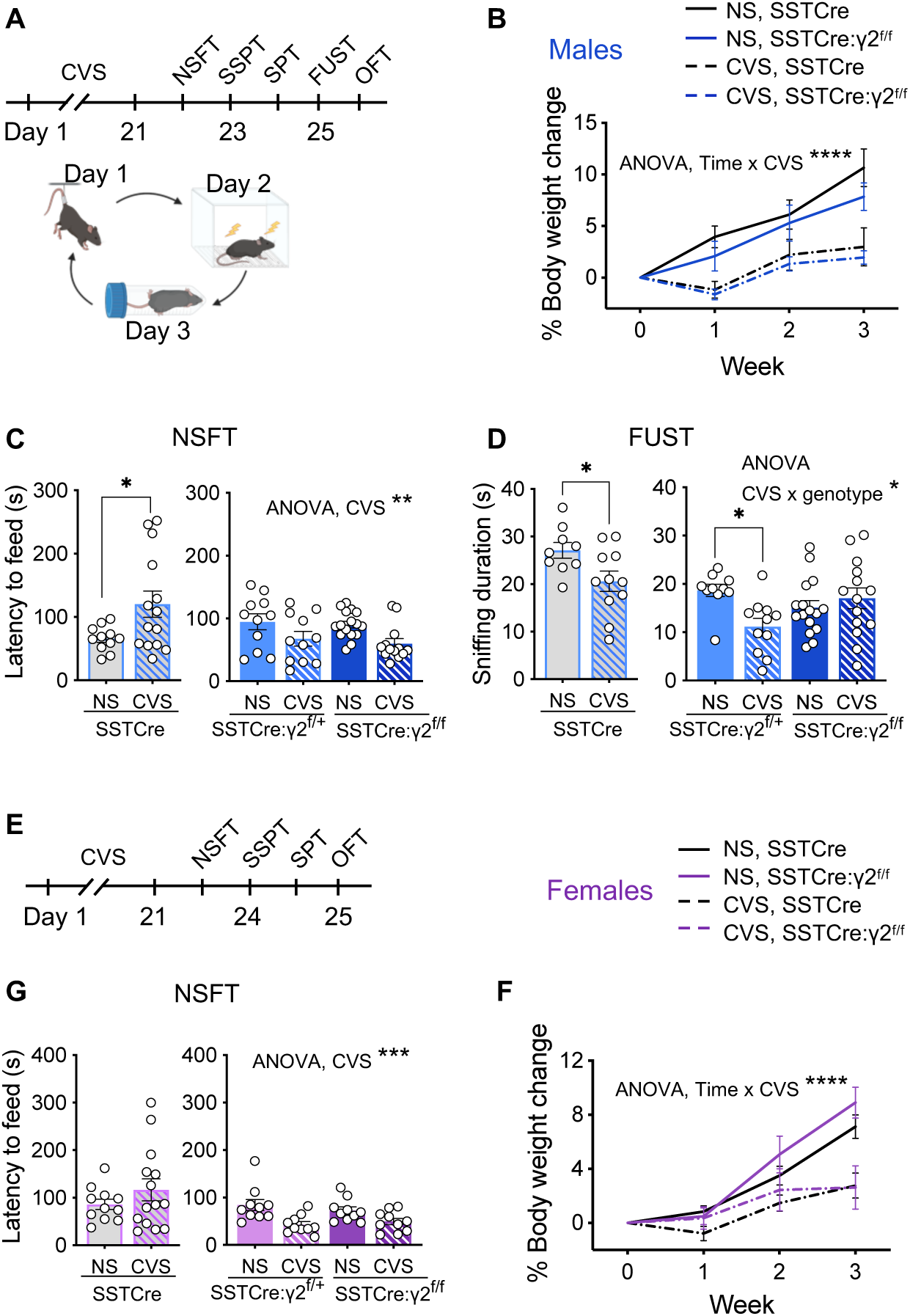
SSTCre:γ2^f/f^ mice are resilient to CVS-induced changes in motivated behavior independent of sex. **A–D)** Data of male mice including time course of experimentation (A) and effects of CVS on body weight changes independent of genotype (B) (CVS effect, F_1, 36_ = 17.08, p < 0.001, time effect, F_1.787, 64.33_ = 47.83, p < 0.0001, time x CVS interaction, F_3, 108_ = 12.94, p < 0.0001, 3-way RM ANOVA). In the NSFT (C), CVS increased the latency to feed in SSTCre mice (p < 0.05, n = 11–14) with opposite effects in SSTCre:γ2^f/+^ and SSTCre:γ2^f/f^ mice (F_1, 47_ = 9.724, p < 0.01). In the FUST (D), CVS reduced the urine sniffing duration of SSTCre mice (p < 0.05, n = 9–11). Comparison of SSTCre:γ2^f/+^ and SSTCre:γ2^f/f^ mice revealed a CVS x genotype interaction (F_1, 47_ = 7.124, p = 0.01) with a CVS-induced reduction in the sniffing time for SSTCre:γ2^f/+^ (p < 0.05, n = 10–11) but not SSTCre:γ2^f/f^ mice. **E–F)** Data of female mice with time course of experimentation (E) and evidence for CVS-induced reductions in body weight independent of genotype (F) (CVS effect, F_1, 35_ = 13.15, p < 0.001, time effect, F_1.86, 65.11_ = 63.08, p < 0.0001, time x CVS interaction, F_3, 105_ = 13.20, p < 0.0001, 3-way RM ANOVA). In the NSFT (G), CVS did not affect the behavior of SSTCre controls but reduced the latency to feed in SSTCre:γ2^f/+^ and SSTCre:γ2^f/f^ mice (F_1, 36_ = 14.73, p < 0.001), similar to males. (C). Bar graphs represent means ± SE. *p < 0.05, **p < 0.01, ***p < 0.001, ****p < 0.0001 (t-test or Tukey’s post hoc test).

In male SSTCre mice, CVS exposure resulted in an increased latency to feed in the NSFT (**Figure 1C**), a decreased time spent sniffing female urine in the FUST (**Figure 1D**), and a reduced time spent in the center in the OFT (**Figure S1A**), indicating both anxiety-like increases in negatively regulated motivated behavior and anhedonia-like reductions in positively regulated motivated behavior. The behavioral measures in the SSPT and SPT were not informative as they were largely unaffected by CVS (**Figure S1B, C**). In striking contrast to SSTCre controls, SSTCre:γ2^f/+^ and SSTCre:γ2^f/f^ mice showed a CVS-induced reduction in the latency to feed in the NSFT, indicating that they were not only resilient to CVS, but that the stress effect was opposite to that observed in SSTCre controls (**Figure 1C**), with the animals seemingly becoming less anxious following stress. In the FUST, SSTCre:γ2^f/f^ but not SSTCre:γ2^f/+^ male mice were resilient to CVS-induced reductions in female urine sniffing duration, indicating that stress resilience extends to positively regulated motivated behavior and that the degree of resilience scales with the level of disinhibition of SST neurons (**Figure 1D**). In the OFT, neither the behavior of SSTCre:γ2^f/+^ mice nor of SSTCre:γ2^f/f^ mice was significantly affected by stress (**Figure S1A**), which is again consistent with stress resilience, even though in this case the mutants did not differ from the SSTCre controls.

Female littermate mice were tested analogously, a couple of weeks after the males for practical reasons. In the NSFT, SSTCre female mice failed to show a CVS effect, in contrast to males. However, CVS of SSTCre:γ2^f/+^ and SSTCre:γ2^f/f^ female mice resulted in a reduced latency to feed (**Figure 1G**), which is indicative of stress resilience similar to males. In the OFT, female SSTCre:γ2^f/+^ and SSTCre:γ2^f/f^ mice showed a CVS effect similar to SSTCre controls (**Figure S1D**). As for males, CVS of female mice had no effect on behavior in the SSPT and SPT (**Figure S1E, F**). In summary, the data suggest that female mice with disinhibited SST neurons are resilient to CVS-induced changes in negatively motivated behavior assessed in the NSFT. Changes in positively motivated behavior could not be assessed as behavior in the SSPT and SPT was unaffected by stress and the FUST is not applicable to females.

### SSTCre:γ2^f/f^ male but not female mice are resilient to CVS-induced changes in the mPFC transcriptome

We next assessed whether disinhibition of SST neurons results in stress resilience at the transcriptome level (**Figure 2A**). We focused on the medial prefrontal cortex (mPFC) as a brain region known to control both positively and negatively regulated forms of motivated behavior. The brains of CVS-exposed mice were harvested 24 h after the last stressor and the mPFC was dissected and processed for RNA-Seq. In male mice, quantitation of CVS-induced DEGs (p < 0.01) from subsamples of SSTCre and SSTCre:γ2^f/f^ mice revealed significantly fewer DEGs in the SSTCre:γ2^f/f^ mice compared to SSTCre controls (**Figure 2B**), indicating that stress resilience is reflected in fewer stress-induced DEGs. Volcano plots of differential expression analyses showed similar numbers of CVS-induced downregulated and upregulated DEGs (p < 0.01) in both genotypes (**Figure 2C, D**). The number of DEGs determined based on all samples of each genotype confirmed the lower number of DEGs in SSTCre:γ2^f/f^ (81) vs SSTCre controls (437). Importantly, heat maps of the DEGs showed that the 437 CVS-induced DEGs observed in the stress-vulnerable SSTCre mice (top row of heat map in **Figure 2C**) were randomly affected by CVS in the stress-resilient SSTCre:γ2^f/f^ mice (bottom row of that heat, note the lack of correspondence in color between the two genotypes). Similarly, the 81 CVS-induced DEGs from the stress-resilient SSTCre:γ2^f/f^ mice (top row of heat map in **Figure 2D**) were randomly affected by CVS in the stress-vulnerable SSTCre mice (bottom row of that heat map). Collectively the data indicate that stress resilience of male SSTCre:γ2^f/f^ mice is reflected in both fewer and qualitatively different CVS-induced DEGs.

**Figure 2.**
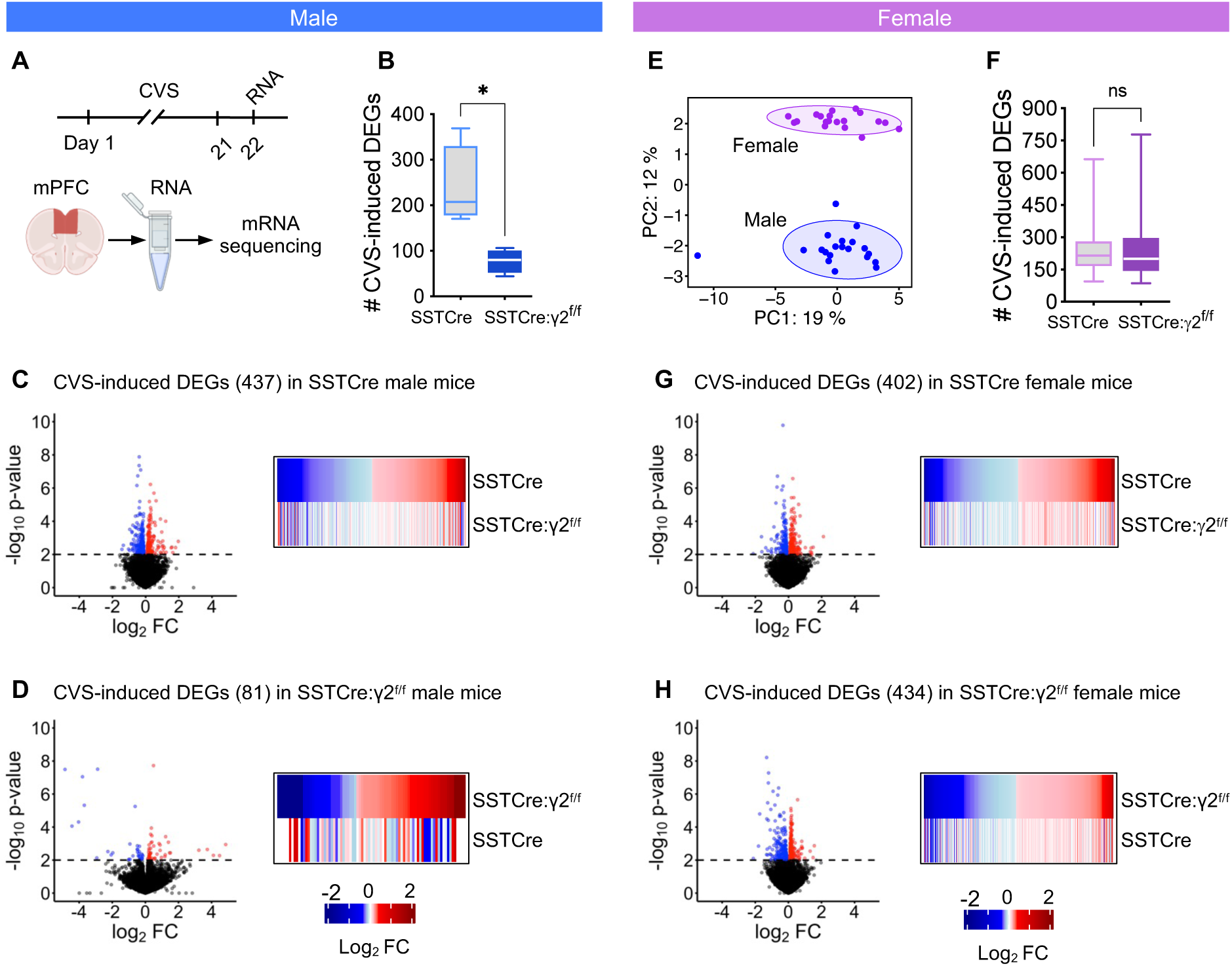
CVS-induced DEGs in the mPFC of SSTCre:γ2^f/f^ stress resilient male mice are fewer and distinct from those of SSTCre controls. **A)** Experimental design. **B)** The average number of CVS induced DEGs (p < 0.01) determined from subsamples of mice revealed fewer CVS-induced DEGs in SSTCre:γ2^f/f^ vs. SSTCre mice (p = 0.013, n = 4, t-test) **C)** Volcano plot of CVS induced DEGs (p < 0.01) in mPFC of SSTCre male mice (n = 5–6), along with heat map comparing the CVS-induced transcriptional changes of DEGs in SSTCre controls (log_2_FC, top) to those of SSTCre:γ2^f/f^ mice. Blue and red dots and lines indicate downregulated and upregulated genes, p value cutoff, respectively. Note that the CVS-induced DEGs of SSTCre mice were randomly affected by CVS in SSTCre:γ2^f/f^ mice. **D)** Volcano plot of CVS induced DEGs (p < 0.01) in SSTCre:γ2^f/f^ males (n = 3–4) along with heat maps comparing transcriptional changes of CVS exposed SSTCre:γ2^f/f^ male mice to those of SSTCre controls. Again, CVS-induced DEGs observed in SSTCre:γ2^f/f^ mice were randomly affected by CVS in SSTCre mice. **E)** PCA analysis revealed clear separation of male and female samples after batch normalization and two outliers among male samples were removed in all analyses. Circles indicate 95% CI. **F–H)** Analyses of female mice analogous to males. The number of CVS-induced DEGs in female mice (p < 0.01) in SSTCre and SSTCre:γ2^f/f^ mice analyzed by subsampling of mice was unaffected by genotype (n = 100, Welch’s t-test) (F). CVS-induced DEGs in the mPFC of female SSTCre mice show comparable directional changes of expression in SSTCre:γ2^f/f^ mice (n = 5) (G) and the CVS-induced DEGs in SSTCre:γ2^f/f^ mice are similarly affected by CVS in SSTCre controls (H).

We next repeated the same analyses for female mice. A comparison of batch normalized male and female transcriptomes by PCA revealed an overt separation of male and female samples. By contrast, the CVS and NS samples of the SSTCre and SSTCre:γ2^f/f^ mice of each sex were co-clustered (**Figure 2E**). Thus, sex differences in the transcriptomes are much larger than the differences induced by CVS and genotype. The quantitation of CVS-induced DEGs from subsamples of female mice revealed similar number of CVS-induced DEGs in the stress-resilient SSTCre:γ2^f/f^ mice compared to the stress-vulnerable SSTCre controls (**Figure 2F**), as is also evident in the volcano blots, showing 402 CVS-induced DEGs across all samples for the stress-vulnerable SSTCre female controls (**Figure 2G**) and 434 CVS-induced DEGs for the SSTCre:γ2^f/f^ stress-resilient female mice (**Figure 2H**). Moreover, the CVS-induced directional gene expression changes of DEGs in the stress-vulnerable SSTCre female mice (top row of heat map in **Figure 2G**) were largely conserved in the stress-resilient SSTCre:γ2^f/f^ female mice (bottom row of heatmap). Similarly, the directional gene expression changes of CVS-induced DEGs seen in the stress-resilient SSTCre:γ2^f/f^ female mice were largely conserved in the stress-vulnerable SSTCre female controls (heatmap in **Figure 2H**). In summary, SSTCre:γ2^f/f^ male mice but not SSTCre:γ2^f/f^ female mice are resilient to CVS-induced changes in the mPFC transcriptome. Therefore, for our further analyses of transcriptomes associated with stress resilience we focused on male mice. We elaborate on sex differences in the brain substrate of stress resilience in the Discussion.

### The CVS-induced transcriptome changes of stress-resilient mice are distinct from the CVS-induced transcriptome changes of stress-vulnerable mice

We argued that putative stress resilience genes should be uniquely affected by CVS in the stress-resilient mice or show opposite CVS effects in the stress-vulnerable compared to stress-resilient mice. A Venn diagram of CVS-induced DEGs of stress-vulnerable (SSTCre) mice and CVS-induced DEGs of stress-resilient (SSTCre:γ2^f/f^) mice revealed 427 CVS-induced DEGs that are uniquely observed in stress-vulnerable mice (for gene lists see **Table S1**), while 71 DEGs were specific for stress-resilient mice (**Figure 3A, Table S2**). A mere 10 CVS-induced DEGs passed the threshold of p < 0.01 in both strains of mice, and nine of these were affected by CVS in the same direction (**Figure 3B**). Notably, Etnk2 showed opposite responses to CVS in stress-vulnerable and stress-resilient mice, which is consistent with a contribution to stress resilience.

**Figure 3.**
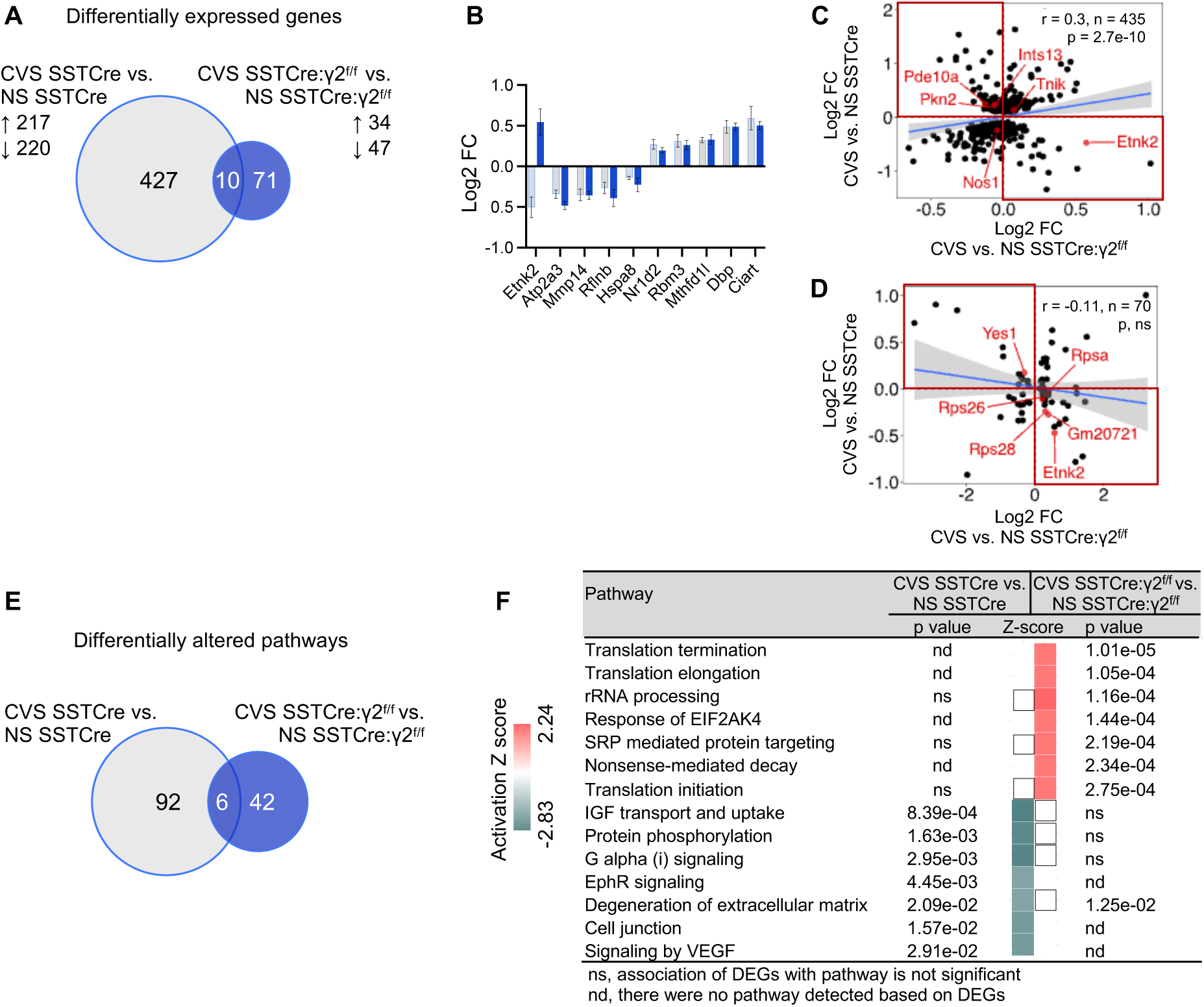
Comparison of CVS-induced DEGs and pathways. A) Venn diagram comparing CVS-induced DEGs (p < 0.01) in SSTCre and SSTCre:γ2^f/f^ mice. The numbers of CVS-induced DEGs in each strain of mice, the numbers of genes affected by CVS in both strains (overlap) and the up- and downregulated genes in each strain are indicated. B) Log2 FC of the 10 DEGs that overlap in (A), ranked based on Log2 FC of CVS SSTCre vs. NS SSTCre from smallest to largest. C, D) Correlational analysis of Log2 FC between CVS-induced DEGs in SSTCre mice and CVS-induced changes of the same genes in SSTCre:γ2^f/f^ mice (C) and between CVS-induced DEGs of SSTCre:γ2^f/f^ mice and CVS induced changes of the same genes in SSTCre mice (D). Note the low correlation in both cases. The top left and bottom right quadrants highlighted in red show DEGs with opposite CVS effects in stress vulnerable vs. stress resilient mice. Grey shades in scatter plots in (C, D) indicate 95% CI. Bar graphs represent means ± SE. E) Venn diagram of IPA pathways (p < 0.05) affected by CVS-induced DEGs in (A). F) Comparison of the top 14 CVS-induced pathways based on absolute Z-scores, ranked by declining p-value and affected significantly (p < 0.05) by at least one of the two contrasts. The top 7 pathways are activated by CVS specifically in stress resilient mice. The next 7 pathways are all inhibited by CVS specifically in stress vulnerable mice. White squares indicate pathways that were detected but a directional Z-score could not be determined. ns, not significant; nd, not detected.

To more comprehensively compare the CVS-induced transcriptome changes of the two strains of mice, we performed a correlational analysis of CVS-induced Log2 FCs of the 437 DEGs observed in the SSTCre mice compared to the Log2 FCs of the same genes in the SSTCre:γ2^f/f^ stress-resilient mice (**Figure 3C**). We then analogously compared the Log2 FCs of the 81 CVS-induced DEGs observed in the SSTCre:γ2^f/f^ stress-resilient mice with the CVS-induced Log2 FCs of the same genes in the SSTCre stress-vulnerable mice (**Figure 3D**). Both of these contrasts revealed negligeable correlation (r = 0.3 and -0.11, respectively), which confirms that the CVS-induced DEGs of the stress-vulnerable and stress-resilient mice are distinct. Putative stress resilience genes include the genes that show opposite CVS effects in the stress-vulnerable vs stress-resilient mice, highlighted by the four red quadrants of **Figure 3C and D (Table S3)**.

### Stress resilience involves chronic stress-induced enhancement of mRNA translation, while stress vulnerability is associated with impairment of diverse signal transduction pathways and reduced translation

To elucidate the function of putative stress resilience genes we performed Ingenuity Pathway analysis (IPA) using DEGs in a default setting. We compared the CVS-induced pathways affected by the 437 CVS-induced DEGs in the stress-vulnerable (SSTCre) mice to those affected by the 81 CVS-induced DEGs of the stress-resilient (SSTCre:γ2^f/f^) mice (**Figure 3E**). IPA revealed 98 CVS-affected pathways (p < 0.05) in stress-vulnerable mice and 48 CVS-affected pathways in stress-resilient mice, with only six pathways affected by CVS in both strains of mice. We then compared the CVS-affected pathways between the two strains using IPA’s integrated comparison analysis tool. The top 14 CVS-regulated pathways (first ranked by Z-score after elimination of pathways related to coronavirus pathogenesis, cancer, autism and pancreatic secretion, and then ranked by p value) revealed a striking segregation of CVS-induced pathways between stress-vulnerable and stress-resilient mice. The seven pathways with the highest Z-scores were all selectively activated in the stress-resilient mice but not in stress-vulnerable mice (**Figure 3F**). The next seven pathways were all selectively inhibited by CVS in the stress-vulnerable mice but not stress-resilient mice. The pathways activated by CVS in stress-resilient mice were all related to mRNA translation and ribosomal RNA processing, and they were principally driven by CVS-induced expression of the same four ribosomal proteins (RPS26, RPS28, RPS29, RPSA) (**Figure 3F, Figure S2**). The pathways inhibited by CVS in stress-vulnerable mice were broadly related to inter- and intracellular signal transduction and cell adhesion.

In an attempt to further corroborate these findings, we performed pathway analyses of the 180 genes that showed opposite CVS effects in stress-resilient (SSTCre:γ2^f/f^) vs stress-vulnerable (SSTCre) mice, i.e. the genes that mapped to the four quadrants marked in red in **Figure 3C, D**. These 180 genes were significantly altered by CVS (p < 0.01) in one of the two strains except for Etnk2 which was significantly affected in both strains (**Figure 3B**). Based on these DEGs there were a total of 140 pathways that were differentially affected by CVS in the two strains of mice. The top 10 among these pathways ranked by Z-scores included the same translation- and rRNA processing-related pathways activated by CVS in stress-resilient mice (**Figure S3**), confirming that stress resilience is mediated by CVS-induced mRNA translation (for genes underlying pathway changes see **Figure S2**). These same pathways were downregulated by CVS in stress-vulnerable mice (**Figure S3**), although the genes underlying these pathways in this case were only nominally affected by CVS. An additional two pathways related to antigen presentation and gustation were activated in stress-vulnerable mice and inhibited in stress-resilient mice (**Figure S3**). Collectively these data strongly suggest that stress resilience induced by increased activity of SST neurons in the mPFC of male mice involves stress-activated mRNA translation, while stress vulnerability involves stress induced reductions in mRNA translation along with downregulation of diverse inter- and intra-cellular signaling pathways.

A key finding from analyses of the CSDS model was that stress resilience is an active process as evidenced by more numerous stress induced gene expression changes in resilient compared to vulnerable mice and by increased activity of dopaminergic neurons of the ventral tegmental area that project to the nucleus accumbens ^19, 20^. To assess whether corresponding features also apply to the resilience mechanism studied here we used subsampling to compare CVS-induced DEGs of SSTCre:γ2^f/f^ vs NS SSTCre controls (corresponding to stress resilient vs NS control mice of the CSDS model) and of CVS SSTCre vs NS SSTCre controls (corresponding to stress susceptible vs NS control mice of the CSDS model). Indeed, CVS-exposed stress-resilient (SSTCre:γ2^f/f^) mice showed significantly greater number of DEGs than CVS exposed stress-vulnerable (SSTCre) controls (**Figure S4**) Thus, resilience driven by increased activity of SST neurons in the mPFC fits the definition of an ‘active’ process, analogous to that driven by dopaminergic neurons in the reward circuit.

### Stress-resilient mice mimic transcriptomic and pathway changes of stress exposure but without stress axis activation

Acute stress of mice is known to increase the excitability of SST neurons in the mPFC ^14^, which raised the question whether the inverse is also the case: Does increased excitability of SST neurons due to disinhibition of SST neurons mimic the effects of stress exposure? To address this question experimentally we compared the CVS-induced DEGs of stress-vulnerable (SSTCre) mice with the genotype-induced DEGs of stress-resilient vs stress-vulnerable mice (**Figure 4A-D**). A Venn diagram of the two sets of DEGs revealed an overlap of 33 DEGs that showed very similar fold changes with just one gene showing opposite directional effects (**Figure 4B**). Therefore, the DEGs of NS stress-resilient vs stress-vulnerable mice are similarly induced by CVS in stress-vulnerable mice. It stands to reason, therefore, that these DEGs represent putative stress resilience genes that are naturally induced by chronic stress exposure even though these gene expression changes are insufficient to induce stress resilience. A broader correlational analysis of the Log2 FCs of the 437 CVS-induced DEGs in stress-vulnerable mice with the Log2 FCs of the same genes in NS stress-resilient vs stress-vulnerable mice revealed a strong correlation between CVS-induced and genotype-induced gene expression changes (r = 0.78, **Figure 4C**) that was confirmed by a similarly strong correlation of Log2 FCs of the 270 genotype-induced DEGs and the CVS-induced Log2 FCs of the same genes in the stress-vulnerable mice (r = 0.67, **Figure 4D**).

**Figure 4.**
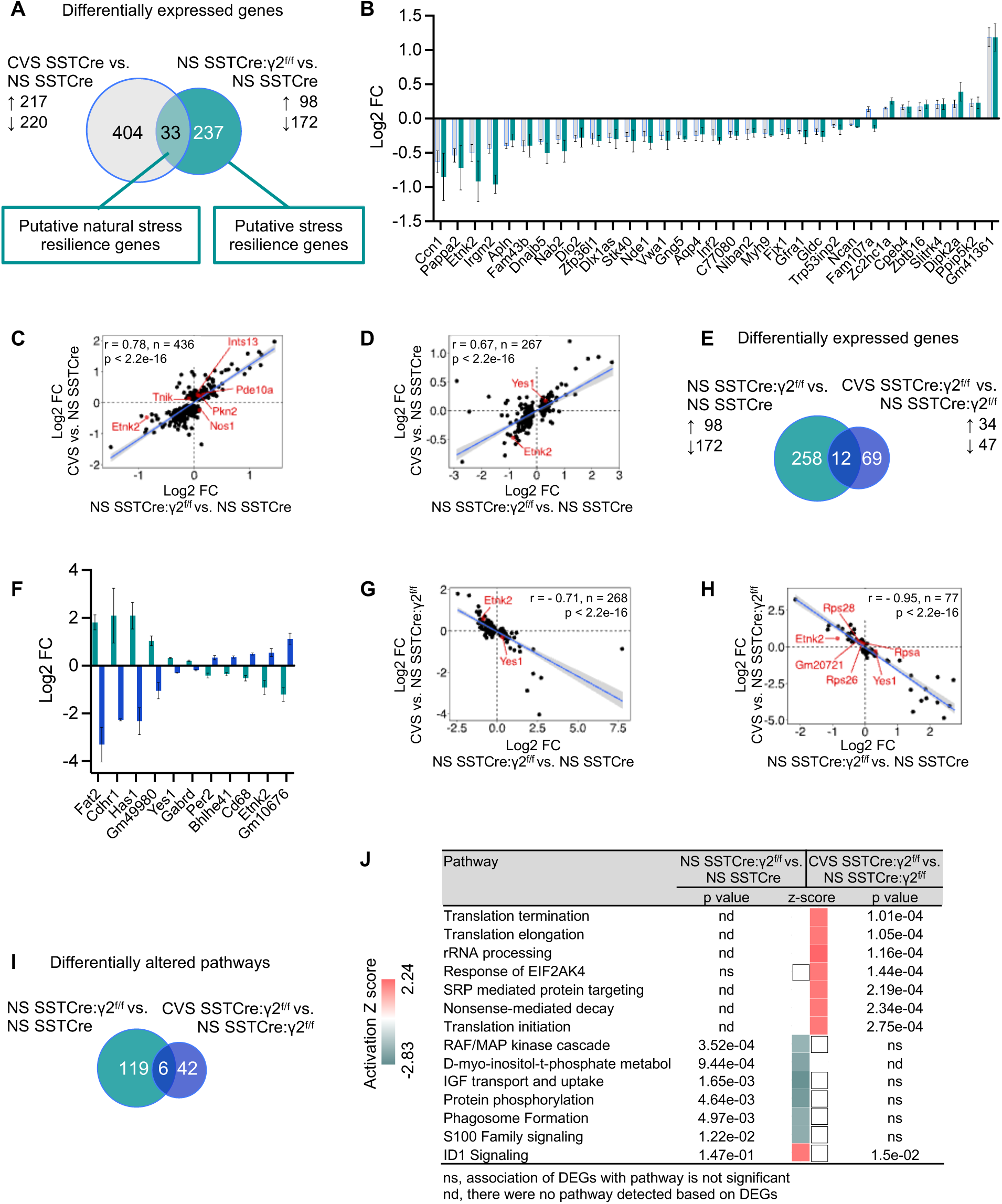
NS stress-resilient male mice mimic transcriptome changes of CVS-exposed stress vulnerable mice, and they are reversed by CVS exposure. **A)** Venn diagram comparing CVS-induced DEGs in SSTCre mice with genotype induced DEGs in NS SSTCre:γ2^f/f^ vs. NS SSTCre mice. **B)** Log2 FC of 33 DEGs that overlap in (A). **C**, **D)** Correlation of Log2 FC between CVS-induced DEGs in SSTCre mice and genotype-induced changes in expression of the same genes in NS SSTCre:γ2^f/f^ vs. NS SSTCre mice (C), and between genotype-induced DEGs (NS SSTCre:γ2^f/f^ vs. NS SSTCre mice) and CVS-induced changes of the same genes in SSTCre mice (D). Note the high level of correlation of FCs in both contrasts. **E)** Venn diagram comparing genotype induced DEGs in NS SSTCre:γ2^f/f^ vs. NS SSTCre mice with CVS-induced DEGs in SSTCre:γ2^f/f^ mice. **F)** Log2 FC of the 12 DEGs that overlap in (E). Note that the genotype induced gene expression change of each is opposite to the CVS effect in the stress resilient mutants. **G, H)** Correlational analyses of Log2 FC of genotype induced DEGs (NS SSTCre:γ2^f/f^ vs. NS SSTCre mice) with CVS-induced changes of the same genes in SSTCre:γ2^f/f^ mice (G) and Log2 FC of CVS-induced DEGs in SSTCre:γ2^f/f^ mice with genotype-induced changes in expression of the same genes in NS SSTCre:γ2^f/f^ vs. NS SSTCre mice (H). Note the strong anticorrelation of FCs between these two contrasts. **I**) Venn diagram of IPA pathways (p < 0.05) induced by CVS in stress-resilient mice compared to pathways induced by genotype (NS SSTCre:γ2^f/f^ vs. NS SSTCre mice). **J**) The top seven pathways are activated selectively by CVS in stress-resilient mice as in Figure 3F. Six of the next seven pathways are selectively inhibited in NS stress-resilient mice. White squares indicate pathways that were detected but a directional Z-score could not be determined. ns, not significant; nd, not detected.

We next compared the pathways affected in NS stress-resilient vs stress-vulnerable mice with those induced by CVS exposure of stress-vulnerable mice. Among the top 15 pathways ranked by Z-score and p values, nine pathways were inhibited under both conditions, and only three pathways had opposing Z-scores (activation under one condition and inhibition in the other) (**Figure S5**). The majority of pathways affected were related to aspects of signal transduction. Therefore, the pathway changes of NS stress-resilient (SSTCre:γ2^f/f^) mice compared to NS stress-vulnerable mice mimic aspects of CVS exposure of stress-vulnerable (SSTCre) mice.

The notion that NS stress-resilient mice show transcriptome changes that are correlated with transcriptome changes of CVS-exposure of stress-vulnerable mice raised the question of whether the stress-resilient mice show evidence of constitutive activation of the hypothalamus-pituitary-adrenal (HPA) axis. To address this possibility, we compared the serum Cort levels of NS and CVS-exposed SSTCre (stress-vulnerable) and SSTCre:γ2^f/f^ (stress-resilient) mice nine days after 21 days of CVS exposure. The stress-resilient (SSTCre:γ2^f/f^) mice showed a strong trend towards reduced serum Cort, independent of prior CVS exposure, and there were no lasting effects of CVS on Cort levels, independent of genotype (**Figure S6**). Therefore, although the transcriptome changes of the stress-resilient mice mimic those of chronic stress exposure of stress-vulnerable mice, stress resilience does not involve constitutive activation of the HPA axis.

### Chronic stress exposure of stress-resilient mice results in reversal of constitutive gene expression changes of stress-resilient mice

One of the above 33 putative natural stress resilience genes, Etnk2, that is downregulated both in NS stress-resilient vs stress-vulnerable mice and in CVS-exposed stress-vulnerable mice (**Figure 4B**) stood out already earlier as a candidate stress resilience gene that is upregulated by stress in stress-resilient mice (**Figure 3B**), indicating that CVS exposure of stress-resilient mice reversed the downregulation of Etnk2 seen in NS stress-resilient vs stress-vulnerable mice. To address whether similar gene expression changes were associated more broadly with stress resilience, we compared the genotype-induced DEGs of stress-resilient vs stress-vulnerable mice with the CVS-induced DEGs of stress-resilient mice (**Figure 4E–H, Table S2, S4**). There were 12 overlapping DEGs (including Etnk2) that passed the significance threshold for both contrasts. They all showed directional changes induced by CVS in stress-resilient mice that were opposite to those induced by genotype in the absence of stress (**Figure 4F**). To further compare the effect sizes of all DEGs of the two contrasts we first compared the Log2 FCs of the 270 genotype-induced DEGs to the Log2 FCs of the same genes induced by CVS in SSTCre:γ2^f/f^ mice. Strikingly, these two factors were almost perfectly anticorrelated (r = -0.71) **(Figure 4G**). Similarly, the Log2 FCs of the 81 CVS-induced DEGs of SSTCre:γ2^f/f^ mice were strongly anticorrelated with the Log2 FCs of the same genes in the genotype comparison (r = - 0.95) **(Figure 4H**). Thus, CVS exposure of the stress-resilient mice results in reversal of the baseline/constitutive transcriptome changes of the stress-resilient mice.

Comparison of the pathways induced by the above DEGs confirmed that CVS exposure of stress-resilient mice involved strong activation of pathways related to mRNA translation as seen earlier (**Figure 4I, J** compared to **Figure 3F**). Six of the next seven pathways were selectively inhibited in NS stress-resilient vs stress-vulnerable mice. As also noted earlier, the pathways inhibited in NS stress-resilient vs stress vulnerable mice were largely the same as the ones that were inhibited by CVS of stress-vulnerable mice (**Figure S5).**

### Stress-induced DEGs of stress-vulnerable but not stress-resilient mice are prominently associated with risk genes of human stress-related psychiatric disorders

To examine the relevance of our findings for stress related psychiatric disorders we compared the CVS-induced DEGs of stress-vulnerable and stress-resilient mice with the DisGeNET human gene-disease associations library. Out of 437 CVS-induced DEGs of stress-vulnerable (SSTCre) mice, 75 (17.16%) were present in the DisGeNET library for MDD, BP, PTSD, and/or SCZ (**Figure 5**). By contrast, a mere 6 out of 81 (7.4%) CVS-induced DEGs of the stress-resilient (SSTCre:γ2^f/f^) mice were associated with these disorders, and all these remaining DEGs were implicated in MDD. The greater enrichment of disease-associated CVS-induced DEGs of SSTCre compared to SSTCre:γ2^f/f^ mice suggests that the mechanism underlying stress resilience of SSTCre:γ2^f/f^ mice may have therapeutic utility for human stress-associated mental disorders.

**Figure 5.**
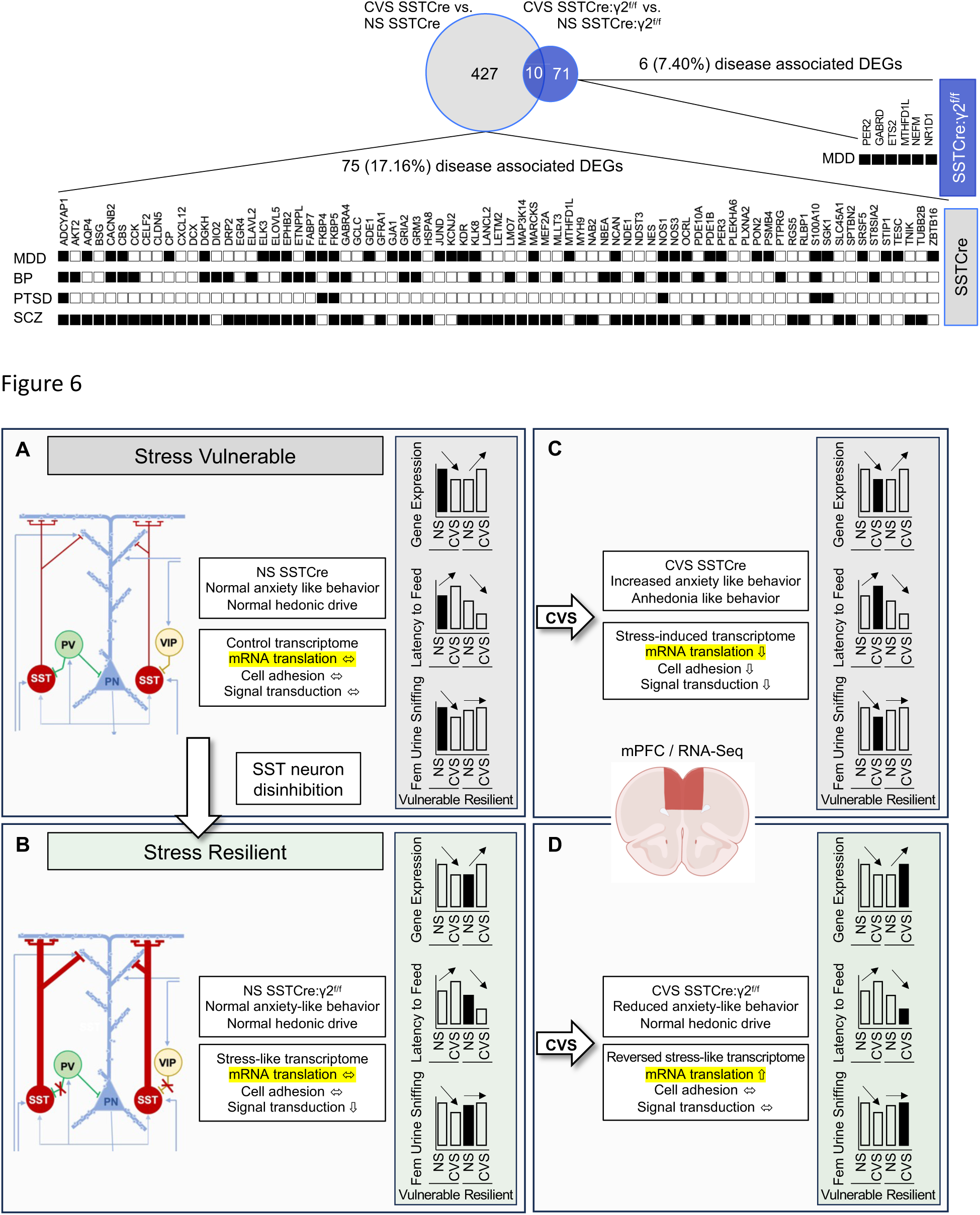
Association of CVS-induced DEGs with human psychiatric disorders. The gene sets of CVS-induced DEGs from male SSTCre (grey) and SSTCre:γ2^f/f^ mice (blue) were searched for genes implicated in human psychiatric disorders in the DisGeNET library using Enrichr. Neurological and psychiatric disorder terms for which the CVS-induced DEGs in SSTCre or SSTCre:γ2^f/f^ mice showed an association (p < 0.05) were selected. Shown are the 75 CVS-induced DEGs in SSTCre male mice associated with MDD (sum of terms Unipolar Depression, Depressive Disorder, Mental Depression and Postpartum Depression, p < 0.05 for all four terms), Bipolar Disorder (BP, p < 0.001), Posttraumatic Stress Disorder (PTSD, p < 0.05) or schizophrenia (SCZ, p < 0.001), respectively. A corresponding analysis of the 81 CVS-induced DEGs in SSTCre:γ2^f/f^ mice revealed 6 DEGs associated with MDD (p < 0.05) and none with the other disorders.

## DISCUSSION

We here have presented a comprehensive transcriptomic analysis of chronic stress resilience of mice, induced by disinhibition of SST-positive GABAergic interneurons. We first show that mice with disinhibited SST neurons are resilient to CVS-induced anxiety and anhedonia-like defects in motivated behavior. Using a milder but longer lasting chronic stress paradigm that was continued throughout behavioral testing we previously reported that stress resilience of these mice was limited to males and to anxiety-like behavior ^8^. However, using CVS as a shorter duration (3-week) but more intense stress protocol we show here that resilience extends to stress induced anhedonia and female mice. Importantly, in the NSFT the mice with disinhibited SST neurons were not only resilient to stress but they showed an inverted stress response, indicating that CVS resulted in an anxiolytic-like reduction of aversion from the test situation. In previous experiments using uncontrolled chronic mild stress as a stressor, the stressed SSTCre:γ2^f/f^ male mice showed unaltered behavior compared to NS controls, suggesting that the behavioral response to chronic stress depends on the nature or intensity of the chronic stressor.

As the next major finding, we report that stress resilience of male mice with disinhibited SST neurons is associated with fewer and distinct CVS-induced DEGs in the mPFC (**Figure 2B-D**), a feature that is reflected in negligible correlation of CVS-induced gene expression changes between stress-vulnerable and stress-resilient mice (**Figure 3C-D**). By contrast, the CVS-induced DEGs of female stress-vulnerable and stress-resilient mice were similar in numbers (**Figure 2F**), and the directional changes in expression of DEGs remained correlated (r = 0.44 and 0.49, respectively, not shown). In a separate study that is currently under review we have used a stereotaxically-targeted chemogenetic approach to map SST neuron mediated stress resilience to specific brain regions ^21^. Selective activation of SST neurons in the mPFC resulted in resilience of male but not female mice while activation of SST neurons in the ventral hippocampus led to resilience in female but not male mice. Our transcriptome studies here confirm that stress resilience in the mPFC is male specific.

As a third major finding we show that SST neuron-mediated stress resilience in the mPFC of male mice is associated with stress-induced enhanced translation, using two types of overlap analyses of DEGs. The DEGs that were affected by CVS in stress-resilient but not stress-vulnerable mice mapped to multiple pathways indicating CVS-enhanced mRNA translation. By contrast, stress vulnerability was associated with CVS-induced downregulation of cell adhesion and signal transduction pathways. A separate pathway analyses of genes that showed opposite CVS-induced changes in transcript levels in stress-resilient vs stress-vulnerable mice again indicated that stress resilience involves enhanced mRNA translation. This second contrast additionally pointed to impaired mRNA translation in stress-vulnerable mice (for a schematic summary see Figure 6). Notably, previous studies have identified endoplasmic reticulum (ER) stress and corresponding impairment of mRNA translation as a cellular mechanism contributing to the detrimental effects of chronic stress exposure ^9^. It seems logical, therefore, that enhanced translation serves as a mechanism that promotes stress resilience, in addition to the prevention of stress-induced downregulation of cell adhesion and signal transduction pathways. However, while chronic stress-induced ER stress has been mapped to SST neurons ^40^, our bulk tissue level transcriptome data necessarily suggest that the stress resilience consequences of SST-neuron-mediated enhanced translation are not limited to this sparse cell type.

**Figure 6.** Graphic summary of results. **A, B**) Schematic of GABAergic microcircuit of the mPFC, including the three major types of GABAergic interneurons in stress-vulnerable (A) and stress-resilient mice (B). The stress-resilient mice (SSTCre:γ2^f/f^ mice) lack postsynaptic GABA_A_ receptors in SST neurons, which results in disinhibition of SST neurons and enhanced GABAergic inhibition mainly at distal apical dendrites of cortical output neurons. **C**) CVS exposure of SSTCre mice results in heightened anxiety and anhedonia-like behavior and stress-induced transcriptome changes in the mPFC, along with pathway changes indicative of reduced signal transduction and nominally reduced mRNA translation. Similar stress induced transcriptome changes are observed in the NS stress-resilient mice (B), along with pathway changes indicating reduced signal transduction. **D**) CVS exposure of stress resilient mice triggers the reversal of stress-like transcriptome changes observed in NS stress-resilient mice, including normalization of signal transduction pathways but results in activation of mRNA translation pathways. Bar graphs in shaded boxes illustrate gene expression changes in the mPFC across the four conditions of a representative putative stress resilience gene (i.e. Etnk2, **Figure S7A**), along with anxiety- and anhedonia-like behavioral changes. Note the opposite, bidirectional CVS-induced changes in gene expression and behavior in stress-vulnerable vs. stress-resilient mice.

As a fourth major finding, we found that stress resilience induced by SST neuron activation in NS stress-resilient mice involves transcriptome changes that mimic chronic stress exposure. A direct comparison of biochemical pathways affected by CVS of stress-vulnerable (SSTCre) mice and by disinhibition of SST neurons in stress-resilient (SSTCre:γ2^f/f^) mice showed that 9 of the 15 most prominently changed signal transduction pathways were inhibited by both conditions (**Figure S5**), with only three pathways affected in opposite directions, which confirms that stress resilience involves pathway changes that mimic chronic stress exposure. Even more striking, chronic stress exposure of NS stress-resilient mice resulted in reversal of the stress-like transcriptome signature along with normalization of pathway changes seen in the NS stress-resilient mice. Importantly, the stress-like transcriptome signature of NS stress-resilient mice was associated with a trend towards lower serum CORT and therefore did not involve activation of the HPA axis. Moreover, the behavior of NS SSTCre:γ2^f/f^ mice was indistinguishable from that of NS SSTCre:γ2^f/+^ littermates, which confirms that the chronic stress-like transcriptome signature of NS stress-resilient mice did not involve systemic or behavioral stress.

Lastly, we found that stress induced DEGs of stress-vulnerable mice show greater association with disease genes of human psychiatric disorders than stress-induced DEGs of stress-resilient mice. This suggests that differences in GABAergic mechanisms underlying stress resilience contribute to vulnerability and resilience to human stress-associated psychiatric disorders.

Putative stress resilience genes highlighted in **Figure 4C, D, G and H** that showed differential expression in stress vulnerable vs stress resilient mice fell into two classes. A first class of DEGs showed a basal change in expression in stress-resilient mice that mimicked the CVS-induced change in expression in stress-vulnerable mice, which was then normalized by CVS exposure of stress-resilient mice (for representative genes see **Figure S7A**). These types of DEGs add to the growing body of evidence that stress resilience is an active process that involves greater or opposite changes in CVS-induced gene expression compared to CVS in stress-vulnerable mice and is not simply due to the absence of or a reduced chronic stress response ^41^. A second, less common class of DEGs was affected less by CVS (or genotype x CVS) in the stress-resilient compared to stress-vulnerable mice (**Figure S7B**). Future work will need to address whether altered expression of any of these DEGs is sufficient to confer resilience to chronic stress exposure.

In conclusion, defects in GABAergic inhibition at dendrites of pyramidal cells are increasingly recognized as a cellular mechanism of vulnerability for stress-associated mental disorders ^11, 40,42^. Conversely, increasing GABAergic inhibition at dendrites promotes resilience through enhanced mRNA translation as shown here for male mice in the mPFC. Future experiments will need to address whether similar mechanisms operate in the ventral hippocampus of female mice. The transcriptomic signature of stress resilience mimics that of stress exposure, suggesting that stress in moderation may come with lasting neuroprotective properties.

## Acknowledgments

We thank Dr. Istvan Albert, Dr. Aswathy Sebastian and Dr. Nicole Lazar for expert advice and Yao Guo for technical assistance. This publication was made possible by a grant (MH099851) from the National Institute of Mental Health (NIMH) to B.L. and generous support from Penn State University. Its contents are solely the responsibility of the authors and do not necessarily represent the views of Penn State or of the NIMH.

## BIBLIOGRAPHY

1. Morris HM, Hashimoto T, Lewis DA. Alterations in somatostatin mRNA expression in the dorsolateral prefrontal cortex of subjects with schizophrenia or schizoaffective disorder. Cereb Cortex 2008; 18(7): 1575–1587.

2. Sibille E, Morris HM, Kota RS, Lewis DA. GABA-related transcripts in the dorsolateral prefrontal cortex in mood disorders. Int J Neuropsychopharmacol 2011; 14(6): 721–734.

3. Tripp A, Kota RS, Lewis DA, Sibille E. Reduced somatostatin in subgenual anterior cingulate cortex in major depression. Neurobiol Dis 2011; 42(1): 116–124.

4. Fee C, Banasr M, Sibille E. Somatostatin-Positive Gamma-Aminobutyric Acid Interneuron Deficits in Depression: Cortical Microcircuit and Therapeutic Perspectives. Biol Psychiatry 2017; 82(8): 549–559.

5. Davies P, Katzman R, Terry RD. Reduced somatostatin-like immunoreactivity in cerebral cortex from cases of Alzheimer disease and Alzheimer senile dementa. Nature 1980; 288(5788): 279–280.

6. Chen Y, Hunter E, Arbabi K, Guet-McCreight A, Consens M, Felsky D et al. Robust differences in cortical cell type proportions across healthy human aging inferred through cross-dataset transcriptome analyses. Neurobiology of aging 2023; 125: 49–61.

7. Mathys H, Peng Z, Boix CA, Victor MB, Leary N, Babu S et al. Single-cell atlas reveals correlates of high cognitive function, dementia, and resilience to Alzheimer’s disease pathology. Cell 2023; 186(20): 4365–4385 e4327.

8. Jefferson SJ, Feng M, Chon U, Guo Y, Kim Y, Luscher B. Disinhibition of somatostatin interneurons confers resilience to stress in male but not female mice. Neurobiol Stress 2020; 13: 100238.

9. Lin LC, Sibille E. Somatostatin, neuronal vulnerability and behavioral emotionality. Mol Psychiatry 2015; 20(3): 377–387.

10. Lyu J, Nagarajan R, Kambali M, Wang M, Rudolph U. Selective inhibition of somatostatin-positive dentate hilar interneurons induces age-related cellular changes and cognitive dysfunction. PNAS Nexus 2023; 2(5): pgad134.

11. Luscher B, Maguire JL, Rudolph U, Sibille E. GABA(A) receptors as targets for treating affective and cognitive symptoms of depression. Trends Pharmacol Sci 2023; 44(9): 586–600.

12. Rudy B, Fishell G, Lee S, Hjerling-Leffler J. Three groups of interneurons account for nearly 100% of neocortical GABAergic neurons. Dev Neurobiol 2011; 71(1): 45–61.

13. Chiu CQ, Lur G, Morse TM, Carnevale NT, Ellis-Davies GC, Higley MJ. Compartmentalization of GABAergic inhibition by dendritic spines. Science 2013; 340(6133): 759–762.

14. Joffe ME, Maksymetz J, Luschinger JR, Dogra S, Ferranti AS, Luessen DJ et al. Acute restraint stress redirects prefrontal cortex circuit function through mGlu(5) receptor plasticity on somatostatin-expressing interneurons. Neuron 2022; 110(6): 1068–1083 e1065.

15. Chiu CQ, Martenson JS, Yamazaki M, Natsume R, Sakimura K, Tomita S et al. Input-Specific NMDAR-Dependent Potentiation of Dendritic GABAergic Inhibition. Neuron 2018; 97(2): 368–377 e363.

16. Fuchs T, Jefferson SJ, Hooper A, Yee PH, Maguire J, Luscher B. Disinhibition of somatostatin-positive GABAergic interneurons results in an anxiolytic and antidepressant-like brain state. Mol Psychiatry 2017; 22(6): 920–930.

17. LaPlant Q, Chakravarty S, Vialou V, Mukherjee S, Koo JW, Kalahasti G et al. Role of nuclear factor kappaB in ovarian hormone-mediated stress hypersensitivity in female mice. Biol Psychiatry 2009; 65(10): 874–880.

18. Labonte B, Engmann O, Purushothaman I, Menard C, Wang J, Tan C et al. Sex-specific transcriptional signatures in human depression. Nat Med 2017; 23(9): 1102–1111.

19. Krishnan V, Han MH, Graham DL, Berton O, Renthal W, Russo SJ et al. Molecular adaptations underlying susceptibility and resistance to social defeat in brain reward regions. Cell 2007; 131(2): 391–404.

20. Nestler EJ, Russo SJ. Neurobiological basis of stress resilience. Neuron 2024; 112(12): 1911–1929.

21. Jiang T, Feng M, Hutson A, Guo Y, Luscher B. Sex-specific GABAergic microcircuits that switch vulnerability into resilience to stress and reverse the effects of chronic stress exposure. biorx 2024. doi: 10.1101/2024.07.09.602716

22. Schweizer C, Balsiger S, Bluethmann H, Mansuy IM, Fritschy JM, Mohler H et al. The gamma 2 subunit of GABA(A) receptors is required for maintenance of receptors at mature synapses. Mol Cell Neurosci 2003; 24(2): 442–450.

23. Whitten WK. Occurence of anoestrus in mice caged in groups. J Endocrinol 1959; 18: 102–107.

24. Shen Q, Lal R, Luellen BA, Earnheart JC, Andrews AM, Luscher B. gamma-Aminobutyric acid-type A receptor deficits cause hypothalamic-pituitary-adrenal axis hyperactivity and antidepressant drug sensitivity reminiscent of melancholic forms of depression. Biol Psychiatry 2010; 68(6): 512–520.

25. Isingrini E, Camus V, Le Guisquet AM, Pingaud M, Devers S, Belzung C. Association between repeated unpredictable chronic mild stress (UCMS) procedures with a high fat diet: a model of fluoxetine resistance in mice. PLoS One 2010; 5(4): e10404.

26. Feng M, Crowley NA, Patel A, Guo Y, Bugni SE, Luscher B. Reversal of a Treatment-Resistant, Depression-Related Brain State with the Kv7 Channel Opener Retigabine. Neuroscience 2019; 406: 109–125.

27. Chen S. Ultrafast one-pass FASTQ data preprocessing, quality control, and deduplication using fastp. iMeta 2023; 2(2).

28. Dobin A, Davis CA, Schlesinger F, Drenkow J, Zaleski C, Jha S et al. STAR: ultrafast universal RNA-seq aligner. Bioinformatics 2013; 29(1): 15–21.

29. Liao Y, Smyth GK, Shi W. featureCounts: an efficient general purpose program for assigning sequence reads to genomic features. Bioinformatics 2014; 30(7): 923–930.

30. Love MI, Huber W, Anders S. Moderated estimation of fold change and dispersion for RNA-seq data with DESeq2. Genome Biol 2014; 15(12): 550.

31. Zhang Y, Parmigiani G, Johnson WE. ComBat-seq: batch effect adjustment for RNA-seq count data. NAR Genom Bioinform 2020; 2(3): lqaa078.

32. Chen EY, Tan CM, Kou Y, Duan Q, Wang Z, Meirelles GV et al. Enrichr: interactive and collaborative HTML5 gene list enrichment analysis tool. BMC Bioinformatics 2013; 14: 128.

33. Kuleshov MV, Jones MR, Rouillard AD, Fernandez NF, Duan Q, Wang Z et al. Enrichr: a comprehensive gene set enrichment analysis web server 2016 update. Nucleic Acids Res 2016; 44(W1): W90–97.

34. Xie Z, Bailey A, Kuleshov MV, Clarke DJB, Evangelista JE, Jenkins SL et al. Gene Set Knowledge Discovery with Enrichr. Curr Protoc 2021; 1(3): e90.

35. Pinero J, Ramirez-Anguita JM, Sauch-Pitarch J, Ronzano F, Centeno E, Sanz F et al. The DisGeNET knowledge platform for disease genomics: 2019 update. Nucleic Acids Res 2020; 48(D1): D845–D855.

36. Pinero J, Bravo A, Queralt-Rosinach N, Gutierrez-Sacristan A, Deu-Pons J, Centeno E et al. DisGeNET: a comprehensive platform integrating information on human disease-associated genes and variants. Nucleic Acids Res 2017; 45(D1): D833–D839.

37. Pinero J, Queralt-Rosinach N, Bravo A, Deu-Pons J, Bauer-Mehren A, Baron M et al. DisGeNET: a discovery platform for the dynamical exploration of human diseases and their genes. Database (Oxford) 2015; 2015: bav028.

38. Pinero J, Sauch J, Sanz F, Furlong LI. The DisGeNET cytoscape app: Exploring and visualizing disease genomics data. Comput Struct Biotechnol J 2021; 19: 2960–2967.

39. Koopmans F, van Nierop P, Andres-Alonso M, Byrnes A, Cijsouw T, Coba MP et al. SynGO: An Evidence-Based, Expert-Curated Knowledge Base for the Synapse. Neuron 2019; 103(2): 217–234 e214.

40. Tomoda T, Sumitomo A, Newton D, Sibille E. Molecular origin of somatostatin-positive neuron vulnerability. Mol Psychiatry 2022; 27(4): 2304–2314.

41. Friedman AK, Walsh JJ, Juarez B, Ku SM, Chaudhury D, Wang J et al. Enhancing depression mechanisms in midbrain dopamine neurons achieves homeostatic resilience. Science 2014; 344(6181): 313–319.

42. Ren Z, Sahir N, Murakami S, Luellen BA, Earnheart JC, Lal R et al. Defects in dendrite and spine maturation and synaptogenesis associated with an anxious-depressive-like phenotype of GABAA receptor-deficient mice. Neuropharmacology 2015; 88: 171–179

