## Supplementary Figures for "Transcriptome signatures of the medial prefrontal cortex underlying GABAergic control of resilience to chronic stress exposure"

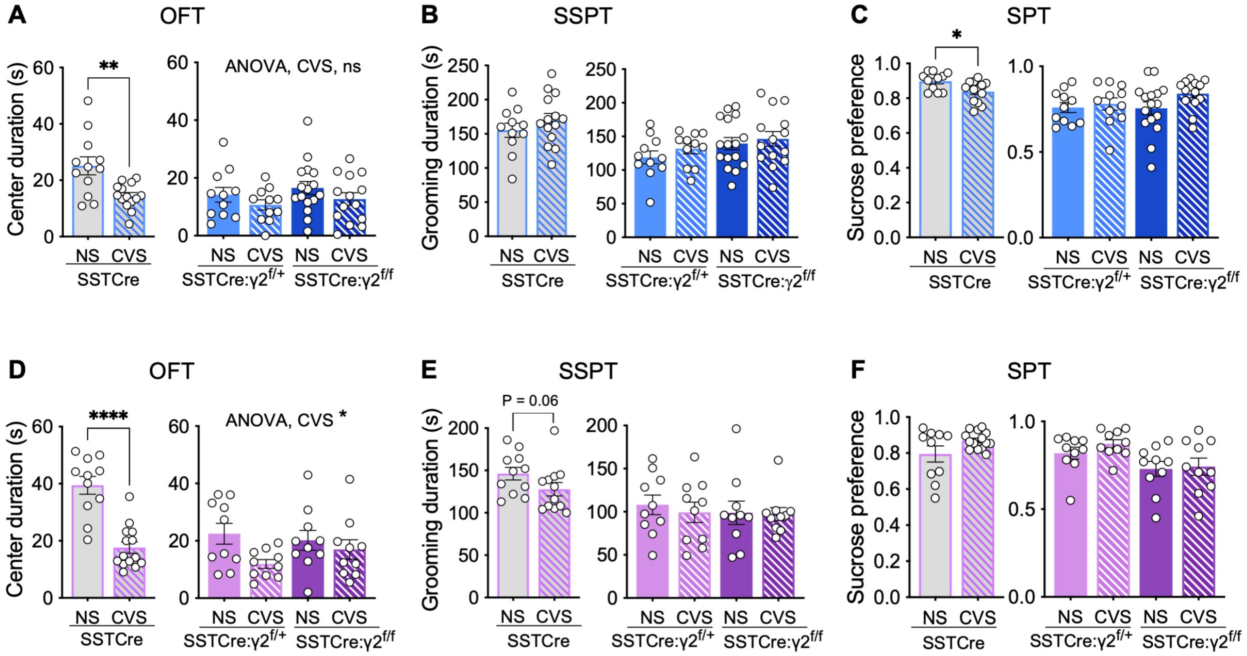


**Figure S1. SSTCre:$\gamma 2^{f/f}$ mice are resilient to CVS-induced behavior independent of sex. A–C)** Male mice. In the OFT (A), CVS reduced the center duration in SSTCre mice ($p < 0.01$, t-test), but not SSTCre:$\gamma 2^{f/+}$ and SSTCre:$\gamma 2^{f/f}$ mice. In the SSPT (B), CVS had no effect on grooming duration independent of genotype. In the SPT (C), CVS reduced the sucrose preference of SSTCre mice ($p < 0.05$, t-test) but not SSTCre:$\gamma 2^{f/+}$ and SSTCre:$\gamma 2^{f/f}$ mice ($F_{1, 48} = 2.56$, p, ns). **D–F)** Female mice. In the OFT (D), CVS reduced the center duration of SSTCre mice ($p < 0.0001$) with less robust effects in the SSTCre:$\gamma 2^{f/+}$ and SSTCre:$\gamma 2^{f/f}$ mutants ($F_{1, 36} = 4.866$, $p < 0.05$). In the SSPT (E), CVS resulted in a trend of a reduction in grooming duration in SSTCre ($p = 0.06$, Mann Whitney test), but not SSTCre:$\gamma 2^{f/+}$ and SSTCre:$\gamma 2^{f/f}$ mice ($F_{1, 36} = 0.2067$, ns). In the SPT (F), CVS had on effect on sucrose preference independent of genotype. Bar graph represent means ± SE. *$p < 0.05$, n = 11–14 for all groups, t-test or Mann Whitney test.

| **CVS SSTCre:γ2$^{f/f}$ vs. NS SSTCre:γ2$^{f/f}$ (Figure 3F, 4J, upregulated pathways)** | | | |
|---|---|---|---|
| Pathways | p value | Z-score | Molecules |
| Translation termination | 1.00e-04 | 2 | Rps26, Rps28, Rps29, Rpsa |
| Translation elongation | 1.05e-04 | 2 | Rps26, Rps28, Rps29, Rpsa |
| rRNA processing | 1.15e-04 | 2.24 | Fbl,Rps26, Rps28, Rps29, Rpsa |
| Response of EIF2AK4 | 1.45e-04 | 2 | Rps26, Rps28, Rps29, Rpsa |
| SRP mediated protein targeting | 2.19e-04 | 2 | Rps26, Rps28, Rps29, Rpsa |
| Nonsense-mediated decay | 2.34e-04 | 2 | Rps26, Rps28, Rps29, Rpsa |
| Translation initiation | 2.75e-04 | 2 | Rps26, Rps28, Rps29, Rpsa |
| **CVS SSTCre vs. NS SSTCre (Figure 3F, downregulated pathways)** | | | |
| Pathways | p value | Z-score | Molecules |
| IGF transport and uptake | 8.32e-04 | -2.83 | Ccn1, Chrdl1, Ckap4, Cp, Lamc1, Nucb1, Pappa2, Vwa1 |
| Protein phosphorylation | 1.62e-03 | -2.65 | Ccn1, Chrdl1, Ckap4, Cp, Lamc1, Nucb1, Vwa1 |
| G alpha (i) signaling | 2.95e-03 | -2.71 | Apln, Cx3Cl1, Cxcl12, Gng12, Gng5, Gpr17, Gpr37L1, Grm3, Rgs11, Rgs5, S1Pr2 |
| EphR signaling | 5.13e-03 | -2 | Akt2, Cxcl12, Efnb1, Epha8, Ephb2, Gng12, Gng5, Map3K14, Map4K4 |
| Degeneration of extracellular matrix | 2.09e-02 | -2 | Adamts1, Bsg, Lamc1, Mmp14 |
| Cell junction | 1.58e-02 | -2.24 | Cd151, Cldn5, Fermt2, Jup, Parvb |
| Signaling by VEGF | 2.88e-02 | -2.24 | Akt2, Jup, Kdr, Mapkapk2, Nos3 |
| **Red quadrants in Figure 3C, D** | | | |
| Pathways | p value | Z-score | Molecules |
| Translation termination | 1.00e-04 | 2 | Rps26, Rps28, Rps29, Rpsa |
| Translation elongation | 1.05e-04 | 2 | Rps26, Rps28, Rps29, Rpsa |
| rRNA processing | 1.15e-04 | 2.24 | Fbl,Rps26, Rps28, Rps29, Rpsa |
| Response of EIF2AK4 | 1.45e-04 | 2 | Rps26, Rps28, Rps29, Rpsa |
| SRP mediated protein targeting | 2.19e-04 | 2 | Rps26, Rps28, Rps29, Rpsa |
| Nonsense-mediated decay | 2.34e-04 | 2 | Rps26, Rps28, Rps29, Rpsa |
| Translation initiation | 2.75e-04 | 2 | Rps26, Rps28, Rps29, Rpsa |
| Class I MHC mediated antigen processing and presentation | 9.93e-01 | 1.63 | Fbxl5, Lmo7, Nedd4L, Psmb4, Wwp1, Zbtb16 |
| Mitotic G2-G2/M phases | 9.88e-01 | -1 | Cep70, Nde1, Psmb4, Tubb2B |
| Gustation Pathway | 9.53e-01 | 2 | Cacnb2, Gabra4, Gabrd, Scn4B |

**Figure S2. Key genes underlying differential pathway activation and inhibition**

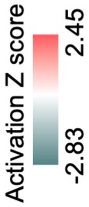


| Pathway | CVS SSTCre vs. NS SSTCre | | CVS SSTCre:$\gamma2^{f/f}$ vs. NS SSTCre:$\gamma2^{f/f}$ | |
|---|---|---|---|---|
| | p value | Z-score | | p value |
| rRNA processing | 8.71e-04 | | | 8.91e-04 |
| Translation termination | 2.40e-03 | | | 2.45e-03 |
| Translation elongation | 2.51e-03 | | | 2.57e-03 |
| Response of EIF2AK4 | 3.31e-03 | | | 3.39e-03 |
| SRP mediated protein targeting | 4.90e-03 | | | 5.01e-03 |
| Nonsense-mediated decay | 5.25e-03 | | | 5.37e-03 |
| Translation initiation | 6.03e-03 | | | 6.17e-03 |
| Class I MHC mediated antigen processing | 2.45e-02 | | | 2.51e-02 |
| Mitotic G2-G2/M phases | 3.09e-02 | | | 3.16e-02 |
| Gustation Pathway | 3.31e-02 | | | 3.39e-02 |

**Figure S3. IPA of genes differentially affected by CVS in SSTCre compared to SSTCre:$\gamma2^{f/f}$ mice.** IPA of the sum of the 180 genes in the four red quadrants of Figure 3C and D that showed opposite CVS effects in stress vulnerable compared to stress resilient mice. Shown are the top 10 pathways affected based on z-scores, displayed from top to bottom in order of increasing p value. Note that nine of the ten most prominently affected pathways are downregulated by CVS in stress vulnerable mice and upregulated by CVS in stress-resilient mice.

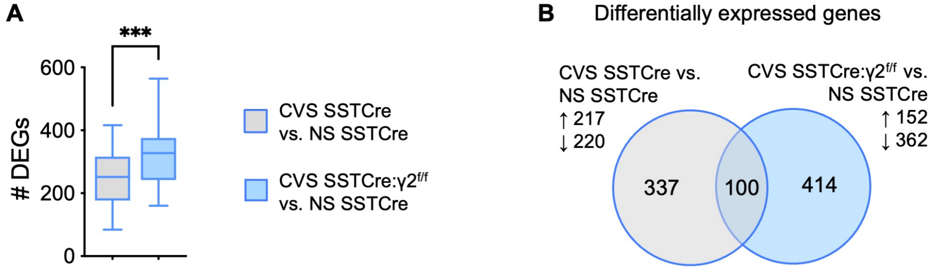


**Figure S4. Comparison of DEGs between CVS exposed SSTCre:$\gamma2^{f/f}$ stress-resilient male mice vs. NS SSTCre controls and CVS exposed SSTCre stress-vulnerable vs. NS SSTCre controls. A)** The average number of CVS plus genotype induced DEGs ($p < 0.01$) in CVS SSTCre:$\gamma2^{f/f}$ vs. NS SSTCre is greater than the DEGs of CVS SSTCre vs. NS SSTCre. **B**) Venn diagram of DEGs ***$p < 0.001$, t-test.

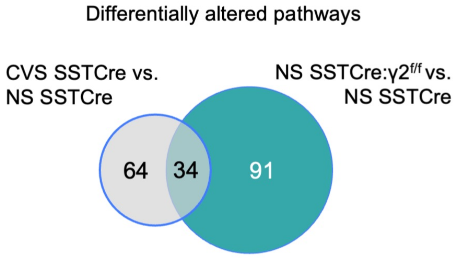


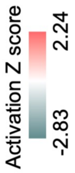


| Pathway | CVS SSTCre vs. NS SSTCre | | NS SSTCre:γ2$^{f/f}$ vs. NS SSTCre | |
|---|---|---|---|---|
| | p value | Z-score | | p value |
| GPCR signaling | 9.12e-03 | | | 3.39e-05 |
| RAF/MAP kinase cascade | ns | | | 3.55e-04 |
| Eicosanoid Signaling | ns | | | 5.89e-04 |
| Inositol phosphate pathway | ns | | | 6.92e-04 |
| EphR signaling | 5.13e-03 | | | 1.07e-03 |
| IGF transport and uptake | 8.32e-04 | | | 1.66e-03 |
| DHA signaling | ns | | | 4.07e-03 |
| Protein phosphorylation | 1.62e-03 | | | 4.68e-03 |
| Phagosome Formation | ns | | | 5.01e-03 |
| S100 Family signaling | ns | | | 1.23e-02 |
| G alpha (i) signaling | 2.95e-03 | | | 4.79e-02 |
| Cell junction organization | 1.58e-02 | | | nd |
| Signaling by VEGF | 2.88e-02 | | | nd |
| Neurovascular coupling signaling | 3.24e-02 | | | 2.08e-01 |
| Macrophage alternative activation | ns | | | 4.37e-02 |

ns, association of DEGs with pathway is not significant
nd, there were no pathway detected based on DEGs

**Figure S5. Venn diagram illustrating overlap between pathways induced by CVS in SSTCre stress-vulnerable mice and disinhibition of SST neurons in the absence of stress [NS SSTCre:γ2$^{f/f}$ vs. NS SSTCre mice].** Note that most pathways are inhibited under both conditions. White squares indicate pathways that were detected but a directional Z-score could not be determined. ns, not significant; nd, not detected.

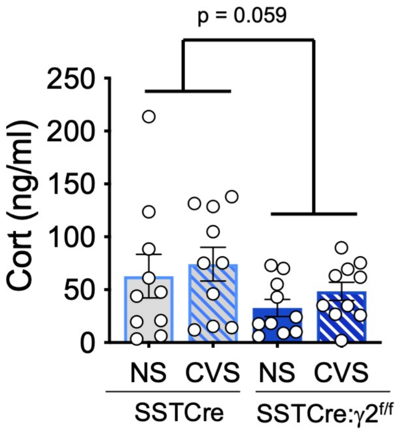


**Figure S6. Analyses of serum corticosterone.** The serum of CVS-exposed, NS SSTCre and SSTCre:γ2$^{f/f}$ male mice harvested 9 days after the end of CVS was subjected to Cort measurements by ELISA. The Cort levels trended lower in stress resilient mice compared to SSTCre controls ($F_{1, 36} = 3.81$, $p = 0.059$) . Bar graphs represent means ± SE.

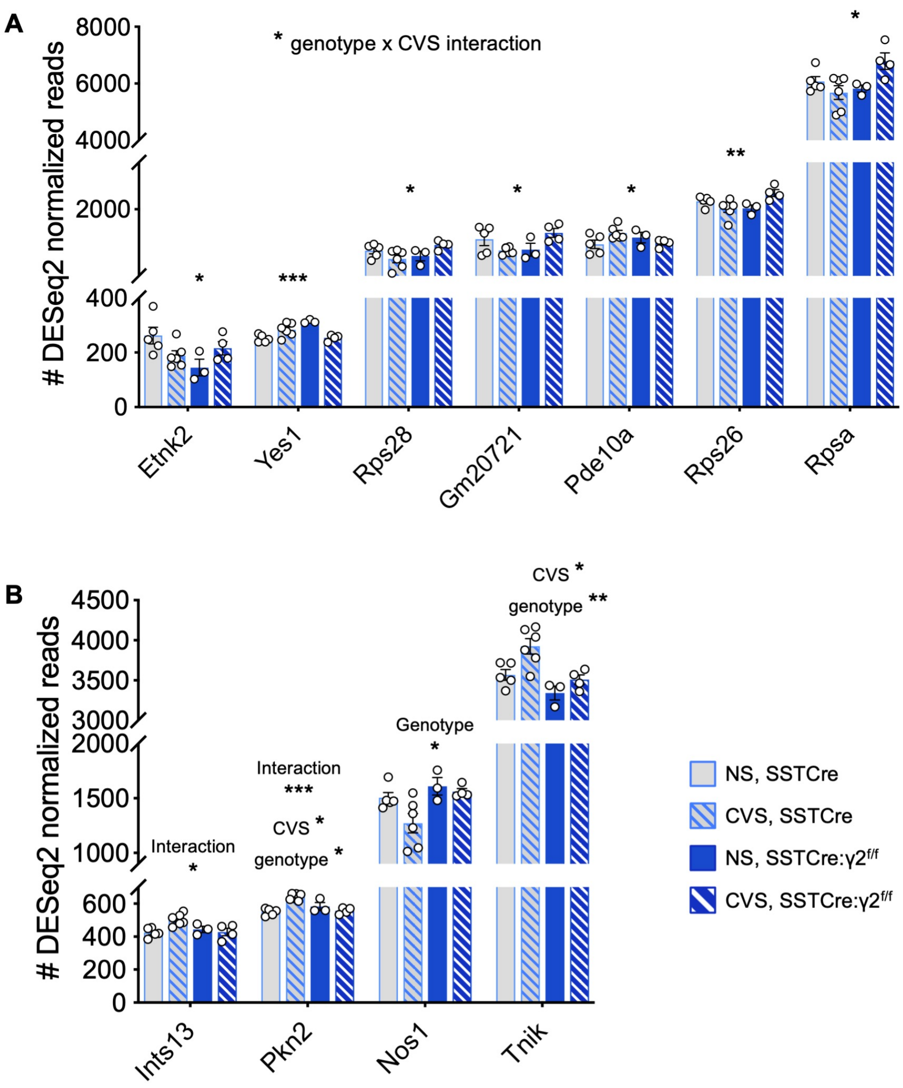


**Figure S7. Two mRNA expression patterns of putative stress-resilient genes. A)** Representative genes that show similar changes in expression in CVS SSTCre and NS SSTCre:γ2$^{f/f}$ compared to NS SSTCre mice that is then normalized by CVS in SSTCre:γ2$^{f/f}$ mice towards NS SSTCre levels, 2-way ANOVAs showed a genotype x CVS interaction for all these genes: Etnk2 ($F_{1, 14} = 7.521$, $p = 0.0159$), Yes1 ($F_{1, 14} = 27.89$, $p = 0.0001$), Rps28 ($F_{1, 14} = 6.508$, $p = 0.0231$), Gm20721 ($F_{1, 14} = 8.840$, $p = 0.0101$), Pde10a ($F_{1, 14} = 5.254$, $p = 0.0379$), Rps26 ($F_{1, 14} = 10.44$, $p = 0.0060$), Rpsa ($F_{1, 14} = 7.783$, $p = 0.0145$). **B)** Representative genes that show smaller CVS effects in SSTCre:γ2$^{f/f}$ vs. SSTCre mice. Inst13, showed a genotype x CVS interaction ($F_{1, 14} = 7.395$, $p = 0.0166$). Genotype, CVS and interaction effects were found for Pkn2 (Genotype, $F_{1, 14} = 4.654$, $p = 0.0489$; CVS, $F_{1, 14} = 7.764$, $p = 0.0146$; interaction, $F_{1, 14} = 24.5$, $p = 0.0002$). Nos1 showed a genotype effect ($F_{1, 14} = 7.055$, $p = 0.0188$). Tnik showed CVS ($F_{1, 14} = 8.777$, $p = 0.0103$) and genotype effects ($F_{1, 14} = 13.29$, $p = 0.0026$; CVS,). *$p < 0.05$, **$p < 0.01$, ***$p < 0.001$.
